# A robust mean and variance test with application to high-dimensional phenotypes

**DOI:** 10.1101/2020.02.06.926584

**Authors:** James R Staley, Frank Windmeijer, Matthew Suderman, Matthew S Lyon, George Davey Smith, Kate Tilling

## Abstract

Most studies of high-dimensional phenotypes focus on assessing differences in mean levels (location) of the phenotype by exposure, e.g. epigenome-wide association studies of DNA methylation at CpG sites. However, identifying effects on the variability (scale) of these outcomes, and combining tests of mean and variability (location-and-scale), could provide additional insights into biological mechanisms. Here, we review variability tests, specifically an extended (for continuous exposures) version of the Brown-Forsythe test, and develop a novel joint location-and-scale score test for both categorical and continuous exposures (JLSsc). The Brown-Forsythe test and JLSsc performed well in comparison to alternative approaches in simulations. These approaches identified >7500 CpG sites that were associated with either mean or variability with gender or gestational age in cord blood methylation in ARIES (Accessible Resource for Integrated Studies). The Brown-Forsythe test and JLSsc are robust tests that can be used to detect associations not solely driven by a mean effect.

## Introduction

Most investigations into health-related phenotypes have focused on determining whether an exposure affects the mean of a phenotype (location test). However, assessing whether an exposure affects the variability of a phenotype (scale test) could also provide insight into the biological mechanisms that control phenotypic variation and disease pathogenesis as well as identify possible interactions^1, 2,^. Furthermore, the potential of combining a location test with a scale test (joint location-and-scale test) has yet to be fully explored, especially in the context of high-dimensional phenotypes where these tests could be used to improve power as well as to identify markers involved in interactions^4^. One example where these approaches could be particularly useful is for epigenome-wide association studies (EWAS), where DNA methylation at CpG (cytosine followed by a guanine) sites across the genome are tested for association with an exposure (Supplementary Text)^5, 6^.

A range of statistical tests have been developed to test whether an exposure affects variability of an outcome, specifically in the context of evaluating variability differences for a continuous variable between groups of individuals^7^. Li *et al*.^8^ compared approaches for assessing methylation variability in the EWAS setting, and showed that the Brown-Forsythe test^9^ performed well compared to alternative approaches. Since this test can be re-formulated in a regression framework^10, 11^, it can be extended to continuous exposures. Methods for jointly testing mean and variability have also been proposed^4, 10, 11, 12, 13, 14, 15^, although these approaches are either limited by sensitivity to distributional assumptions or are restricted to binary exposures.

Here, we review variability tests, specifically the Brown-Forsythe test, and develop a novel joint location-and-scale test, which can be used for both continuous and categorical exposures. We performed a simulation study to compare these approaches to alternative tests, and then applied these modelling approaches to investigate the effect of gender and gestational age on cord blood DNA methylation mean and variability.

## Results

### Review of statistical methods

Ordinary least squares (OLS) regression is commonly used to assess mean differences in an outcome by an exposure and is known to be relatively robust to its underlying assumptions (Methods and Supplementary Text), so is used as the location test. Likewise, the Brown-Forsythe approach has been found to be robust to non-normality and outliers^8^, so is also a good candidate for analysing high-dimensional phenotypes, especially since it can be extended for use with continuous exposures (see Methods). A computationally efficient joint location-and-scale test, which combines *p*-values from location and scale tests using Fisher’s method (JLSp), has been developed for genetic data^10, 11^. However, this test like most existing joint location-and-scale tests requires the phenotype to be normally distributed, while other tests are only designed for binary exposures. Here, we develop a novel joint location-and-scale score test (JLSsc) that is: (i) robust to distributional assumptions, and (ii) can be used for both categorical and continuous exposures.

Our approach essentially combines a location test and scale test, while accounting for the correlation between these tests (see Methods). Briefly, we propose to test the joint null hypothesis *H*_0_: *β* = *δ* = 0 in the model specification:

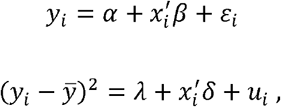

where *y*_*i*_ is the outcome for the *i*-th individual, *x*_*i*_ is the (*k*_*x*_) vector of exposures for the *i*-th individual and 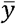 is the sample average of *y*_*i*_.

Let 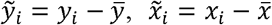 (and 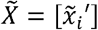) and 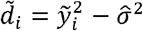, where 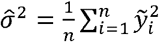

Further, let 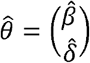, where, and 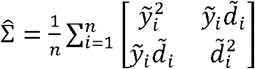, then the score test for *H*_0_: *β* = *δ* = 0, or *H*_0_: *θ* = 0 is given by

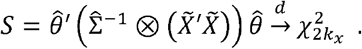

### Simulation study

We performed a simulation study to evaluate the performance of the location and scale tests as well as the joint location-and-scale tests with both binary and continuous exposures in the context of DNA methylation. We compared the Brown-Forsythe scale test to Bartlett’s test^16^ (for simulations with a binary exposure) and a likelihood ratio test (LRT) comparing mixed models with and without a variability effect (LRTv). JLSsc was compared to JLSp as well as a LRT comparing mixed models with and without including both a mean and variability effect for the exposure (LRTmv) and a double generalized linear mixed model (DGLM)^13, 14, 17^. The simulations were performed using data from the Tsaprouni *et al*. study^18^, which investigated the relationship between smoking and DNA methylation (data accessible at NCBI GEO database^19^, accession GSE50660). Type I error simulations were performed across all CpG sites in Tsaprouni *et al*. for varying sample sizes, while power simulations were performed for various phenotypic distributions using 1000 samples where in each simulation replicate the characteristics from a randomly selected CpG site from Tsaprouni *et al*. were used (see Methods and Supplementary Text). In the type I error simulations, common data transformations such as the M-value transformation for DNA methylation data^20^ and inverse normal rank transformation were applied to the methylation data for those approaches that failed to control type I error due to deviations from normality.

OLS regression test of mean differences was not inflated under the null of no mean or variability effect even in 100 samples (Figures 1a and S1). Similarly, the Brown-Forsythe variability test accurately controlled type I error rates (Figures 1b and S2). Bartlett’s test and LRTv had extreme type I error inflation due to the deviations from normality and the existence of outlying values in methylation levels (Figure S3). Likewise, the test statistics from the likelihood-based approaches for joint testing the mean and variability (LRTmv and DGLM) were also heavily inflated (Figure S3). The extreme inflated type I error rates of these approaches were still present after transforming methylation levels using the M-value transformation (Figure S4) but were no longer present after using an inverse normal rank transformation (Figure S5). However, when using this transformation a mean effect can induce a variability effect and vice versa (Figure S6), as seen previously^21^. JLSp fared better than the aforementioned joint tests in controlling type I error rates, although the non-independence of the *p*-values did lead to a small amount of type I error inflation (Figure 1c and Figure S7). The JLSsc approach, on the other hand, correctly controlled type I error rates (Figure 1d and Figure S7).

**Figure 1:**
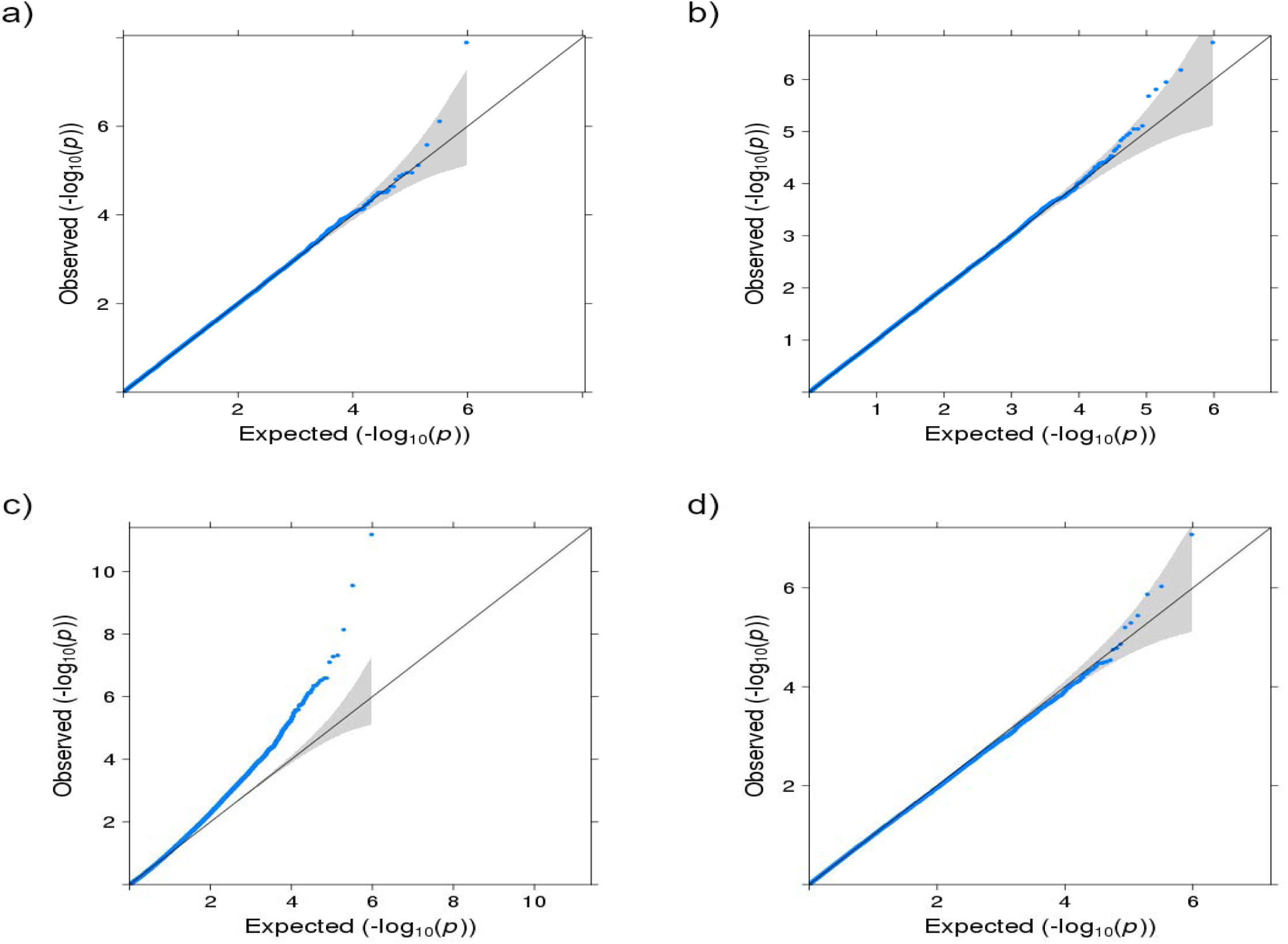
QQ plots for type I error simulations using a binary exposure and 1000 samples. a) OLS (mean test); b) Brown-Forsythe (variability test); c) JLSp (joint test); and d) JLSsc (joint test).

In the power simulations, when there was either a mean or variability effect and the underlying data were normally distributed, the Brown-Forsythe test and JLSsc were less powerful but still performed well in comparison to the equivalent LRT and the alternative approaches (Figure 2). This is expected as the Brown-Forsythe test and JLSsc sacrifice a small amount of power under the normal model for robustness to deviations from this model. Broadly similar results were found when the residual error was heavy-tailed or skewed, when the exposure was a categorical variable with three categories, when a squared exposure term was added to JLSsc for a continuous exposure (although outlier removal in the outcome is necessary here to retain type I error levels), and when there was an outlier in the dataset (Figures S8-S12).

**Figure 2:**
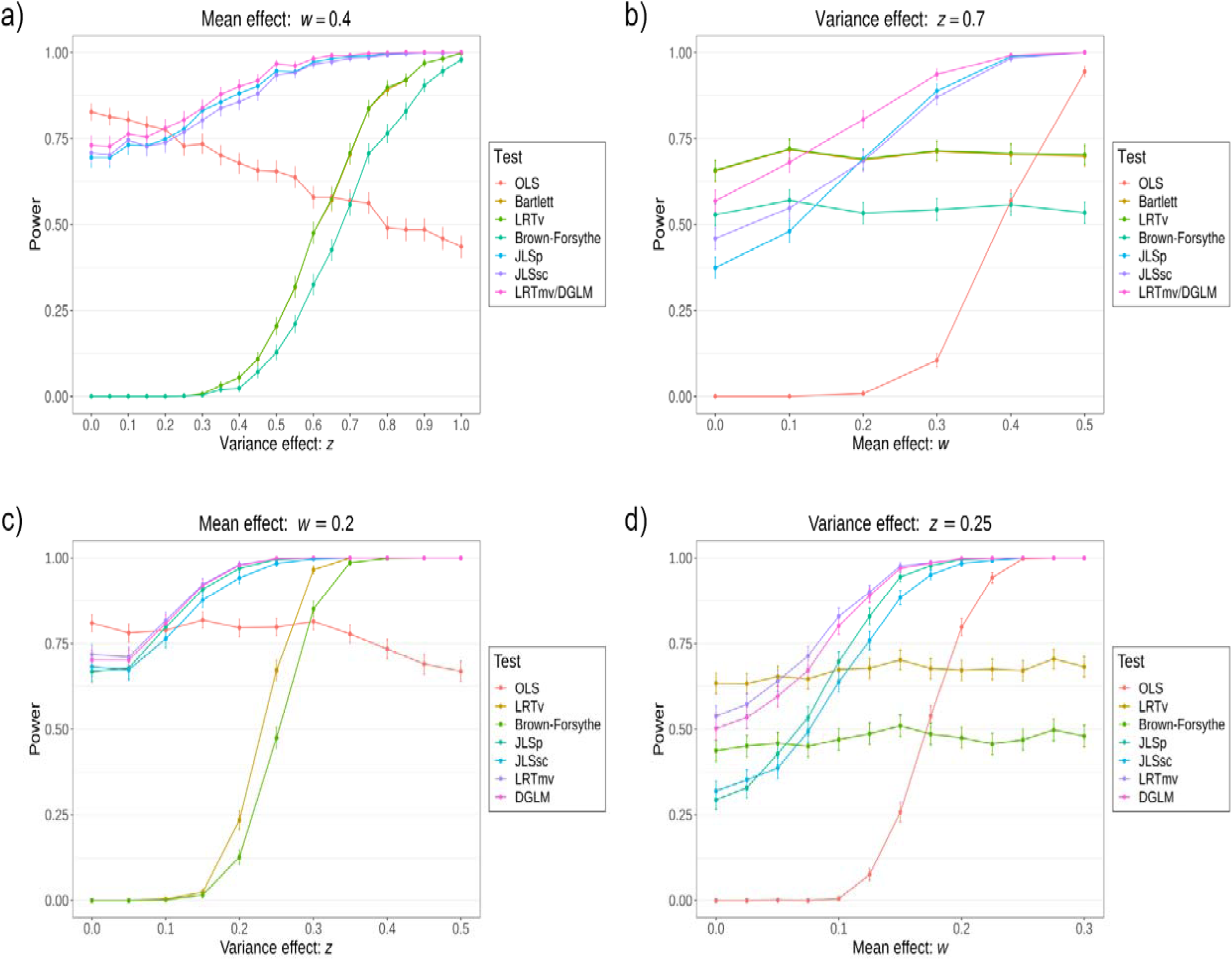
Power simulation results comparing approaches for identifying CpG sites associated with either a mean and/or a variance effect with the exposure at. a) & b) are plots for a binary exposure and c) & d) are plots for a continuous exposure.

The computational time required to complete each approach for 100,000 CpGs with a binary exposure were as follows: 22 minutes for the extended Brown-Forsythe test, 113 minutes for LRTv, 16 minutes for JLSsc and 123 minutes for LRTmv. The relative computation times between the respective variability and joint tests were even greater when the exposure was continuous.

### Application to offspring gender and gestational age on cord blood DNA methylation

We applied the methods that demonstrated good performance in the simulation study, namely OLS regression, the Brown-Forsythe test, JLSp and JLSsc, to identify methylation at CpG sites that are associated with offspring gender and gestational age in the Accessible Resource for Integrated Studies (ARIES) project. ARIES is a sub-sample of the Avon Longitudinal Study of Parents and Children (ALSPAC) (see Methods). In ARIES, 858 children (417 male and 441 female) were available for the analysis of gender, and after excluding offspring with missing maternal information we were left with 708 children (345 males and 363 females) for the analysis of gestational age (mean: 39.5 weeks, standard deviation: 1.5 weeks; Table S1).

Methylation at 8174 CpG sites were associated with gender in cord blood (through the mean, variability or joint tests; Figure 3a and Table S2). Most of these sites were identified through a mean difference in methylation of males and females (7642 CpGs had a mean difference with *p* < 1×10^7^), although 240 CpG sites were associated with a variability difference between males and females. For instance, cg18918831 was more variable in males compared to females (Figure 4). The joint location-and-scale tests identified 7724 of these CpG sites (JLSp identified 7213 sites and JLSsc identified 7228 sites), including all of those with a variability effect. Mean methylation at 5359 of these sites were associated with gender in previous EWAS (Table S2)^22, 23, 24, 25^.

**Figure 3:**
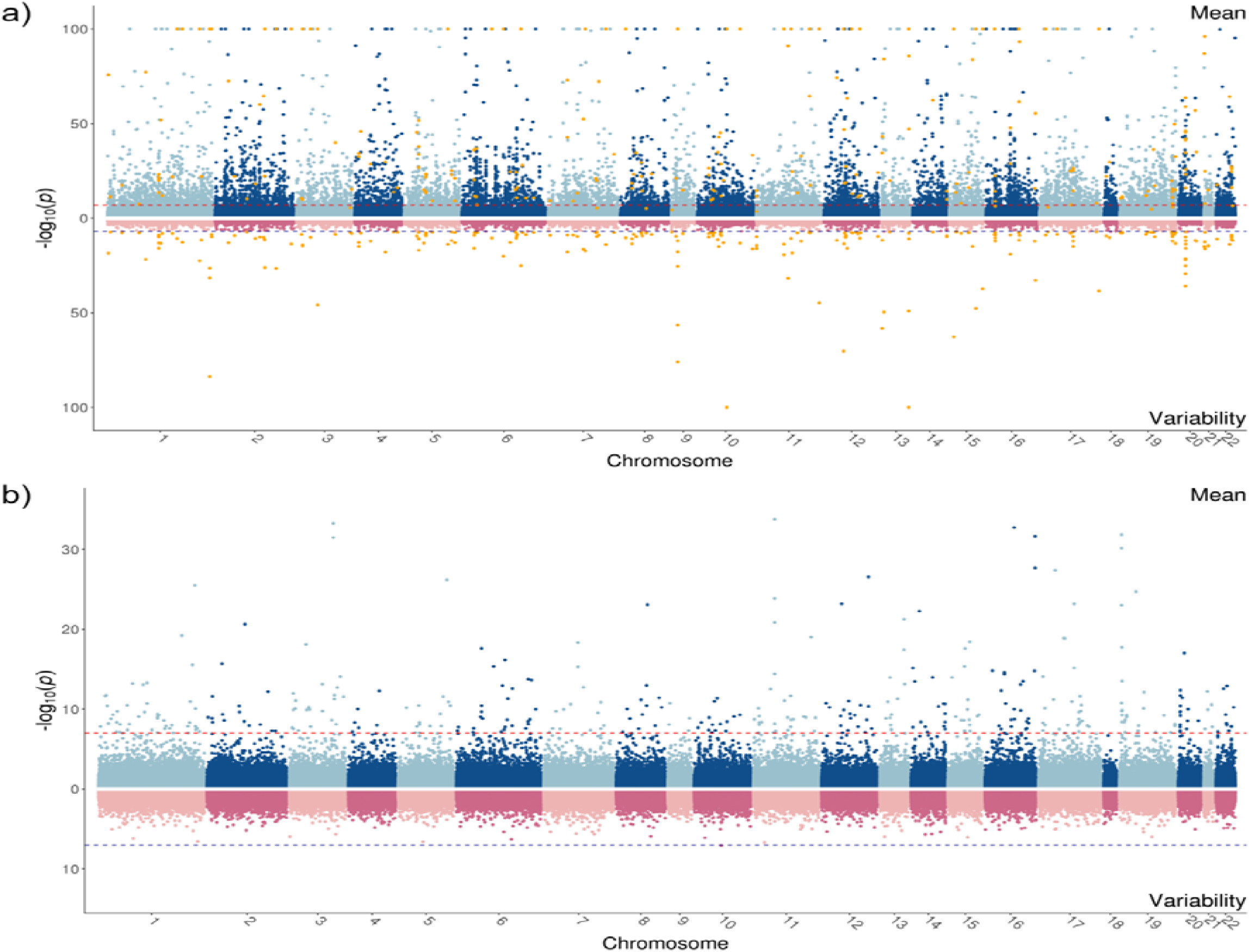
Miami plots for the mean (OLS) and variability (Brown-Forsythe test) associations of methylation with gender a) and gestational age b). The dark red and blue lines represent the threshold and the orange points are CpG sites that are associated with a variance effect.

**Figure 4:**
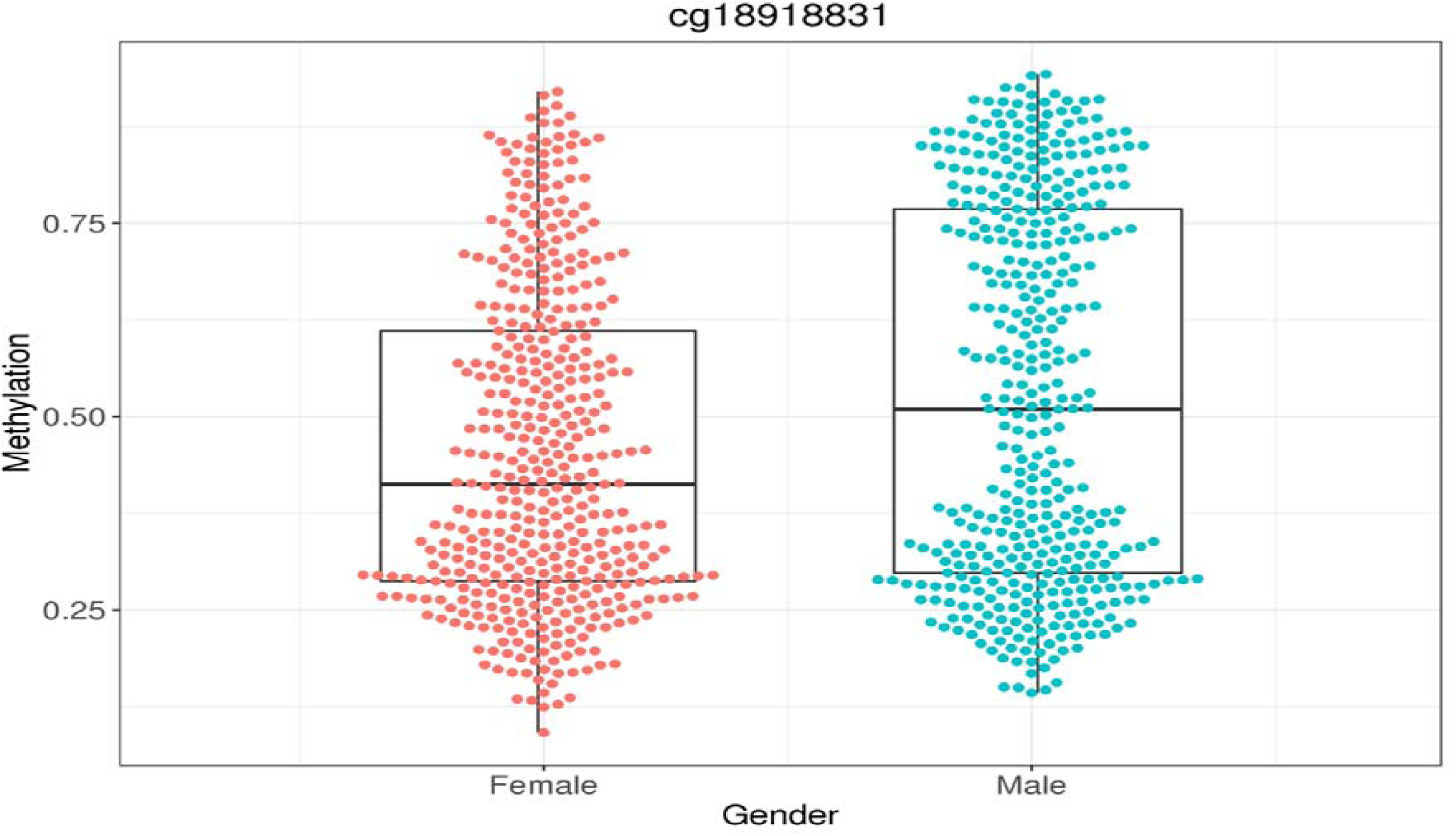
Methylation variability plot for gender.

Gestational age was associated with cord blood methylation at 412 CpG sites (Figure 3b and Table S3). Most of these CpG sites were associated with a mean effect of gestational age on methylation, and there were no CpG sites with a variability effect with *p* < 1x 10^7^. The joint mean and variability tests identified 93.7% of the CpG site associations (JLSp identified 317 and JLSsc identified 340 CpG sites, respectively), including sites that were mostly identified through a variability association (e.g. cg24577594; Table S3). The majority of the CpG sites identified have been found previously in EWAS of gestational age (402 CpG sites; Table S3)^25, 26^.

## Discussion

In this study, we have introduced a framework for testing variability using an extended version of the Brown-Forsythe test and for jointly testing mean and variability. These approaches were compared to the LRTs as well as other alternative methods in simulations and were used to investigate the effect of gender and gestational age on cord blood DNA methylation.

Without transforming the phenotype to be normally distributed, the approaches which assume normality of the phenotype (Bartlett’s test, LRTv, LRTmv and DGLM) had extremely inflated type I error rates when faced with real methylation data. Indeed, these approaches essentially became tests of deviations from normality and outlying values, which can have some utility in identifying outliers caused by disease^27^. However, because of these drawbacks these approaches are not useful for assessing variability nor joint mean and variability effects, especially as normalizing outcome levels to overcome this problem can induce effects that were not present prior to the transformation^21^. The extended Brown-Forsythe test and the JLSsc approach retained correct type I error rates and performed well in comparison to the other approaches in detecting variability and joint effects. These tests were also at least 5 times more efficient than their LRT counterparts.

Over 8000 CpG sites were associated with gender in cord blood methylation, while methylation at 412 CpG sites were associated with gestational age. The majority of these CpG sites were associated with effects of gender and gestational age on mean methylation. However, 240 CpG sites were associated with differences in variability between males and females. JLSsc identified most of the associations in both analyses, except where there was little evidence of a mean/variability effect in the presence of a borderline effect of the other.

These methods are applicable to any area of medical research where variability and joint effects are of interest, although they will be particularly useful for analysing high-dimensional phenotypes where it is not possible to assess the distribution at all markers. For instance, there has been recent interest in using variability tests to attempt to identify gene-environment interactions, as these interactions will often cause heterogeneity in the variance across genotypes^4, 21^. The Brown-Forsythe test has been proposed as a useful test in this scenario^21^, although the extended version presented here and elsewhere^10, 11^ could be used to assess variability trends across genotypes. Furthermore, JLSsc avoids the distributional assumptions made by current methods proposed in the genetics literature^4, 10, 11^.

The limitations of this study also warrant consideration. In the simulations and the applied example, we only analysed DNA methylation data, although we fully expect these results to be generalisable to all phenotypes. The application of the approaches to detect CpG sites associated with gender and gestational age also have several limitations, especially with regards to residual confounding. In particular, there are likely to be other important maternal factors involved in gestation period that we have not adjusted for in our analysis. The ARIES cohort is also not selected at random from the full ALSPAC cohort^28^, and as such, the results from this study may not generalise to the full ALSPAC cohort or the general population.

In summary, the extended Brown-Forsythe test and JLSsc are robust tests of variability and joint mean and variability effects, respectively. These tests can be used in analyses to detect associations for any type of exposure with high-dimensional phenotypes.

## Methods

### Modelling approaches

#### Location tests

Ordinary least squares (OLS) regression is commonly used to assess mean differences in methylation by an exposure. That is,

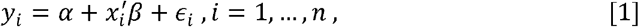

where *y*_*i*_ is the outcome for the *i*-th individual (e.g. DNA methylation levels in epigenome-wide association studies), *x*_*i*_ is the exposure(s) for the -th individual and 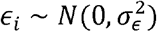 regression is known to be relatively robust to the underlying assumptions related to the residuals when estimating the regression coefficients (discussed further in the Supplementary Text).

#### Scale tests

There are several statistical tests for assessing variability differences of continuous outcome by a categorical exposure^7^. Bartlett’s test^16^ is perhaps the most well-known of these tests (Supplementary Text) and has been used to analyse high-dimensional phenotypes^3, 29^. However, this test is known to be very sensitive to outliers and non-normality of the outcome, which is a major cause of concern when analysing data like DNA methylation. The Brown-Forsythe test^9^, on the other hand, is robust to non-normality of the outcome and outliers^8^. This test is essentially a one-way analysis of variability of the variable 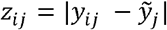, where *y*_*ij*_ is the methylation of the *i*-th individual in the *j*-th group and 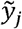 is the median of the *j*-th group. Let 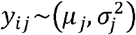, where *μ*_*j*_ and 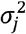 are the mean and variance of *y*_*ij*_ in the *j*-th group, then the test statistic for 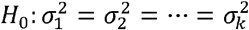 is given by

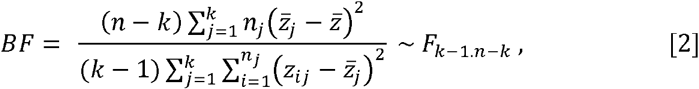

where *k* is the number of groups,*n*_*j*_ is the number of individuals in the *j*-th group and 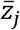 and 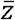 are the group mean and overall mean of *z*_*i j*_, respectively.

The Brown-Forsythe test can be re-formulated as a two-stage approach^10, 11^:

i. Obtain the absolute values of the residuals from a least absolute deviation regression, 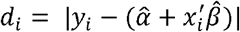.
ii. Test for an association between the *x*_*i*_’s and a function of the *d*_*i*_’s using a regression *F*-test.

Since this regression framework does not depend on the exposure (*x*_*i*_) being categorical, it can also be applied to continuous exposures. Indeed, this approach has the same structure as the Glejser and Bresuch-Pagan tests of heteroskedasticity^30, 31^.

#### Joint location-and-scale tests

If the data are symmetrically distributed then the *p*-values from the location and scale tests are independent and can be combined using Fisher’s method (JLSp)^10, 11^. However, often high-dimensional phenotypes are not all symmetrically distributed (e.g. DNA methylation at CpG sites), which will likely lead to correlated *p*-values for at least some markers. Other alternative approaches for jointly testing for mean and variability effects include likelihood-ratio tests (LRT) comparing linear mixed models with and without including a fixed-effect and random-effect for the exposure (LRTmv) and double generalized linear mixed models (DGLM)^13, 14, 17^ (further details in Supplementary Text). However, these tests are also sensitive to deviations from normality and outlying values^13^.

To alleviate some of the issues involved in testing for mean and variability effects simultaneously, we have developed a joint location-and-scale score test (JLSsc). This approach essentially combines a location test and scale test, while accounting for the correlation between these tests. We propose to test the joint null hypothesis *H*_0_: *β* = *δ* = 0 in the model specification:

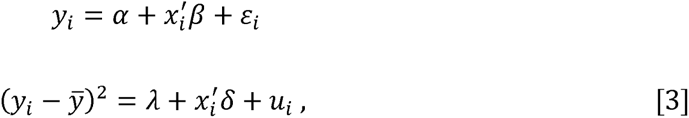

where *x*_*i*_ is a (*k*_*x*_) vector of exposures and 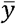 is the sample average of *y*_*i*_. The first part, *H*_0_:*β* = 0, is the null hypothesis that *x* does not affect the mean of *y*. The second part, *H*_0_: *δ* = 0, is the null hypothesis that *x* does not affect the variability of *y*.

Let 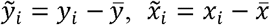 and 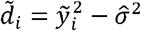, where 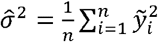. Further, let the *n* x *k*_*x*_ matrix 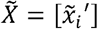 and the *n* vectors 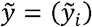 and 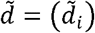 Then the OLS estimators for *β* and *δ* are given by

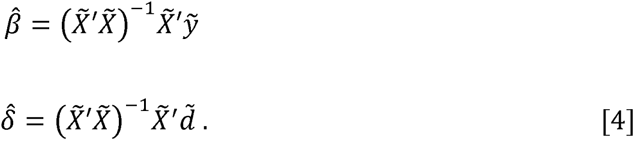

Let 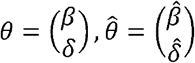 and 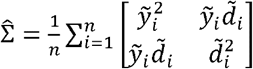. The estimator for the variance of 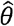 under the null that *β* = *δ* = 0 and the additional assumption that the conditional skewness and kurtosis of *y*_*i*_ do not vary with the values of *x*_*i*_, is then given by

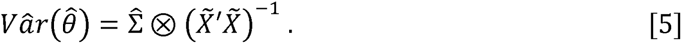

Hence, the score test for HO:, H_0_: *β* = *δ* = 0 or *θ* = 0, is given by

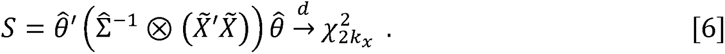

Additional terms such as the square of a continuous exposure, especially useful for modelling the relationship with outcome variability, can be added as part of *x*_*i*_ vector and would be included in both parts of the test. Covariates are regressed out of both the outcome and exposure variables by taking residuals from OLS regression prior to analysis with JLSsc. Further details of JLSsc are discussed in the Supplementary Text.

We have developed an R package to perform these tests available at: https://github.com/jrs95/jlst.

### Simulation study

We assessed the performance of the location and scale tests as well as the joint location-and-scale tests with both binary and continuous exposures in a simulation study based on methylation data. We assessed the performance of OLS regression, Bartlett’s test (for simulations with a binary exposure), Brown-Forsythe test, LRT comparing mixed models with and without a variability effect (LRTv), JLSsc, JLSp, LRTmv and DGLM. For approaches which failed to adequately control type I error rates, we repeated the tests after applying M-value (i.e. log_2_(*y*_*i*_/1-*y*_*i*_)))^20^ and inverse normal rank transformations to the methylation levels. This simulation study was performed based on data from the Tsaprouni *et al*. study^18^, which investigated the relationship between smoking and DNA methylation (data accessible at NCBI GEO database^19^, accession GSE50660).

Type I error simulations were performed by randomly generating a binary or continuous exposure (uncorrelated with mean or variability of any of the methylation levels) and testing the associations across all CpG sites in Tsaprouni *et al*.. To generate datasets with varying sample size (100, 500, 1000 and 10,000 samples), samples were randomly sampled with replacement from the Tsaprouni *et al*. dataset (Supplementary Text). The binary and categorical exposures were randomly generated using *Ber* (0.5)and *N*(0,1), respectively. Quantile-quantile (QQ) plots were used to assess deviations from normality and detect outlying test statistics.

Power simulations were performed using the same exposure distributions as above and setting these exposures to affect the mean and variability of methylation. In each simulation replicate, one CpG was selected at random from the Tsaprouni *et al*. dataset, the mean and standard deviation of this CpG site were used to set the average methylation and to generate mean and variability effects (Supplementary Text). The mean and variability effects of the exposure on methylation were simulated using normal distributions, while the residual error was simulated to be either normally distributed, heavy-tailed or skewed (Supplementary Text). We also performed simulations for a categorical exposure with three categories, adding a squared exposure term to JLSsc for a continuous exposure, and where we generated an outlying value (Supplementary Text). Statistical power was calculated as the proportion of simulation replicates where either the location, scale or joint test had *p* < 1x 10^7^. For each simulation scenario, 1000 simulation replicates were performed for a sample size of 1000 samples.

The computational time of the extended Brown-Forsythe test and JLSsc were compared to their equivalent LRTs for 100,000 randomly selected CpGs from the Tsaprouni *et al*. dataset for the binary and continuous exposures describe above. This analysis was performed using one core (2.6 GHz; 4GB) on a linux server.

### Application to offspring gender and gestational age on cord blood DNA methylation

#### Study population

This study used DNA methylation data generated as part of the Avon Longitudinal Study of Parents and Children (ALSPAC)^32, 33^. ALSPAC recruited 14,541 pregnant women with expected delivery dates between April 1991 and December 1992. Of these initial pregnancies, there were 14,062 live births and 13,988 children who were alive at 1 year of age. Please note that the study website contains details of all the data that is available through a fully searchable data dictionary and variable search tool (http://www.bristol.ac.uk/alspac/researchers/our-data/). Ethical approval for the study was obtained from the ALSPAC Ethics and Law Committee and the Local Research Ethics Committees. Informed consent for the use of data collected via questionnaires and clinics was obtained from participants following the recommendations of the ALSPAC Ethics and Law Committee at the time. Consent for biological samples has been collected in accordance with the Human Tissue Act (2004).

As part of the Accessible Resource for Integrated Studies (ARIES) project (http://www.ariesepigenomics.org.uk)^28^, a sub-sample of 1018 ALSPAC child–mother pairs had DNA methylation measured. The ARIES participants were selected based on availability of DNA samples at two time-points for the mother (antenatal and at follow-up when the offspring was in adolescence) and at three time-points for the offspring (neonatal from cord blood, childhood (age 7) and adolescence (age 17)).

#### Laboratory methods, quality control and pre-processing

The laboratory methods and quality control procedures used have been described elsewhere^34^. In brief, the DNA methylation wet laboratory and pre-processing analyses were performed at the University of Bristol as part of the ARIES project, where the Infinium HumanMethylation450 BeadChip^35^ was used to measure genome-wide DNA methylation levels at over 485,000 CpG sites. The methylation level at each CpG site was calculated as a beta value: the ratio of the methylated probe intensity and the overall intensity. These beta values range from 0 (no methylation) to 1 (complete methylation). The samples were processed using functional normalization with the meffil package^36, 37^. Further quality control procedures are described in the Supplementary Text.

#### Statistical analysis

To investigate the mean and variability effects of gender and gestational age (in weeks, Supplementary Text) on cord blood methylation, we used the approaches which controlled type I error rates without transforming methylation levels, namely OLS regression, the Brown-Forsythe test, JLSp and JLSsc. All analyses were adjusted for cell counts estimated using the method described by de Goede *et al*. for cord blood methylation^38^. We further adjusted for 20 surrogate variables to account for residual batch effects^39^. The gestational age analysis was further adjusted for offspring gender and whether the birth was by caesarean section as well as for maternal characteristics: age, smoking, pre-pregnancy BMI and weight, parity, education, family social class and alcohol intake during pregnancy. CpGs were considered to be associated with either gender or gestational age if one of the location, scale or joint tests had *p* < 1x 10^7^.

All analyses were performed using R (version 3.5.2).

## Acknowledgements

We are extremely grateful to all the families who took part in this study, the midwives for their help in recruiting them, and the whole ALSPAC team, which includes interviewers, computer and laboratory technicians, clerical workers, research scientists, volunteers, managers, receptionists and nurses.

## Author contributions

J.R.S. and M.S. performed the analyses. F.W. and J.R.S. developed the methodology. All authors contributed to the conception of the study and writing the manuscript.

