## Supplementary Text for "A robust mean and variance test with application to high-dimensional phenotypes"

James R Staley^1^, Frank Windmeijer^1,2^, Matthew Suderman^1^, Matthew S Lyon^1,3^, George Davey Smith^1^ and Kate Tilling^1,*^.

^1^MRC Integrative Epidemiology Unit, Population Health Sciences, Bristol Medical School, University of Bristol, Bristol, UK.

^2^Department of Economics, University of Bristol, Bristol, UK.

^3^National Institute for Health Research Bristol Biomedical Research Centre, University of Bristol, Oakfield House, Bristol, BS8 2BN, UK.

^*^Corresponding author.

Correspondence:

Prof Kate Tilling

MRC Integrative Epidemiology Unit, Population Health Sciences, Bristol Medical School, University of Bristol, Bristol, BS8 2BN, UK.

### Supplementary Text

#### Epigenome-wide association studies

Epigenome-wide association studies (EWAS) have been used to assess associations of DNA methylation at CpG (cytosine followed by a guanine) sites from across the genome with diseases and traits^1, 2^. DNA methylation is usually quantified between 0 and 1, and represents the proportion of methylated DNA molecules at the CpG site in the measured tissue. The initial analyses involve univariate testing of each CpG site (e.g. on the Illumina HumanMethylation450 array there are ~485,000 CpG sites^3^) to identify DNA methylation that is associated with an exposure and/or a phenotype^4^ while accounting for multiple testing (a Bonferroni corrected *p*-value threshold of $\sim1\times{10}^{-7}$ is often used for studies based on the Illumina HumanMethylation450 array). Methylation at CpG sites is often treated as the outcome (i.e. a possible consequence of the trait)^4^, where mean levels of methylation are regressed against the exposure using a linear regression model. These analyses are usually adjusted for batch effects and other technical covariates^5^, as well as for cell composition^6^ and other potential confounding factors such as age, gender and other characteristics.

#### Modelling approaches

##### Location tests

Ordinary least squares (OLS) regression is commonly used to assess mean differences in methylation by an exposure. That is,

$$y_{i}=\alpha+x_{i}^{'}\beta+\epsilon_{i} , i=1,\ldots,n , [1]$$

where $y_{i}$ is the outcome for the $i$-th individual (usually DNA methylation measurements in EWAS), $x_{i}$ is the exposure(s) for the $i$-th individual and $\epsilon_{i}\sim N(0, \sigma_{\epsilon}^{2})$. OLS regression is known to be relatively robust to the underlying assumptions related to the residuals when estimating the regression coefficients, particularly the estimated regression coefficient related to the exposure(s), i.e. $\hat{\beta}$, as this is unaffected by the residual mean-zero assumption being violated. Moreover, the assumption that the residuals are normally distributed has less impact on the statistical inference of the regression coefficients with increasing sample size, as for large sample sizes ($n$ > ~100) the central limit theorem applies. The main limitation of use of OLS regression in the context of analysing DNA methylation data is that it is affected by outlying methylation values. However, robust regression techniques^7^ have been developed to handle outlying values in the outcome (e.g. least absolute deviation regression and M-estimators), so this issue is not considered further here.

##### Bartlett’s test of variability

Bartlett’s test^8^ tests the null hypothesis that all $k$ group variances are the same against the alternative that at least one pair of variances differ. That is, let $y_{ij}$ be the outcome of the $i$-th individual in the $j$-th group and $y_{ij}\sim\left( \mu_{j},\sigma_{j}^{2} \right)$, where $\mu_{j}$ and $\sigma_{j}^{2}$ are the mean and variance of $y_{ij}$ in the $j$-th group, then the Bartlett’s test statistic for $H_{0}:\sigma_{1}^{2}=\sigma_{2}^{2}=\ldots=\sigma_{k}^{2}$ is given by

$$BT= \frac{\left( n-k \right)\log\left( S_{p}^{2} \right)- \sum_{j=1}^{k} \left( n_{j}-1 \right)\log(S_{j}^{2})}{1+\frac{1}{3(k-1)}\left( \sum_{j=1}^{k} \left( \frac{1}{n_{j}-1} \right)-\frac{1}{n-k} \right)} \sim\chi_{k-1}^{2} , [2]$$

where $n$ is the total number of individuals, $n_{j}$ is the number of individuals in the $j$-th group and $S_{j}^{2}$ and $S_{p}^{2}$ are the group variance and the pooled variance estimates (i.e. $S_{p}^{2}=\frac{1}{n-k}\sum_{j=1}^{k} \left( n_{j}-1 \right)\log\left( S_{j}^{2} \right)$), respectively.

##### Brown-Forsythe test of variability

The Brown-Forsythe test^9^ is essentially a one-way analysis of variability of the variable $z_{ij}=|y_{ij} -\tilde{y}_{j}|$, where $y_{ij}$ is the methylation of the $i$-th individual in the $j$-th group and $\tilde{y}_{j}$ is the median of the $j$-th group. Hence, the Brown-Forsythe test statistic for $H_{0}:\sigma_{1}^{2}=\sigma_{2}^{2}=\ldots=\sigma_{k}^{2}$ is given by

$$BF= \frac{\left( n-k \right)\sum_{j=1}^{k} n_{j}\left( \bar{z}_{j}-\bar{z} \right)^{2}}{(k-1)\sum_{j=1}^{k} \sum_{i=1}^{n_{j}} \left( z_{ij}-\bar{z}_{j} \right)^{2}} \sim F_{k-1.n-k} , [3]$$

where $k$ is the number of groups, $n_{j}$ is the number of individuals in the $j$-th group and $\bar{z}_{j}$ and $\bar{z}$ are the group mean and overall mean of $z_{ij}$, respectively.

##### Likelihood ratio tests

Linear mixed models^10^ can be used to construct a likelihood ratio test (LRT) of the mean and variability or the variability only. Suppose $y_{i}$ is the response variable for the $i$-th sample and $x_{i}$ is either the binary or continuous exposure, then consider the following model (model 1):

$$y_{i}=\tilde{\alpha}+\tilde{\beta}x_{i}+\epsilon_{i} , i=1,\ldots,n , \left[ 4 \right]$$

where

$$\tilde{\alpha}=\alpha_{0}+u_{1i} ,$$

$$\tilde{\beta}=\beta_{0}+u_{2i} \left[ 5 \right]$$

and $u_{i} \sim N(0, \Sigma_{u})$ (where $\Sigma_{u}$is an unstructured covariance matrix) and $\epsilon_{i}\sim N(0, \sigma_{e}^{2})$.

Consider also the model (model 0):

$$y_{i}=\tilde{\alpha}+\epsilon_{i} , i=1,\ldots,n . \left[ 6 \right]$$

Then to test for a mean and/or variability effect the deviance from model 0 minus the deviance from model 1 is tested against $\chi_{2}^{2}$, i.e. $D\left( model 0 \right)-D\left( model 1 \right) \sim\chi_{2}^{2}$. Likewise, to test for a variability effect only, the deviance from model 1 is tested against the deviance from the same model without including the $u_{2i}$ term using $\chi_{1}^{2}$. For continuous exposures this model assesses the relationship of the variance of $y_{i}$ with $x_{i}$ and $x_{i}^{2}$.

Similarly, a LRT can be constructed using the deviances from double generalized linear models^11^ with and without mean and dispersion parameters related to the exposure ($x_{i}$). The difference here is that a log link is used to model the variability. However, for categorical variables these tests are identical and yield the same results as those from the LRT proposed by Cao *et al*.^12^.

##### Joint location-and-scale test using Fisher’s method (JLSp)

If the outcome data are symmetrically distributed then the $p$-values from the location and scale tests are independent and can be combined using Fisher’s method (JLSp)^13, 14^. That is, we can combine the $p$-value from the location test in [1] ($p_{l}$) with the $p$-value from the location test in [3] ($p_{s}$) using the following test statistic:

$$Q=-2\left( \log\left( p_{l} \right)+\log\left( p_{s} \right) \right) \sim\chi_{4}^{2} . \left[ 7 \right]$$

##### Joint location-and-scale score test (JLSsc)

We propose to test the joint null hypothesis $H_{0}:\beta=\delta=0$ in the model specification:

$$y_{i}=\alpha+x_{i}^{'}\beta+\varepsilon_{i}$$

$$\left( y_{i}-\bar{y} \right)^{2}=\lambda+x_{i}^{'}\delta+u_{i} , [8]$$

where $x_{i}$ is a ($k_{x}$) vector of exposures and $\bar{y}$ is the sample average of $y_{i}$. The first part, $H_{0}:\beta=0$, is the null hypothesis that $x$ does not affect the mean of $y$. The second part, $H_{0}:\delta=0$, is the null hypothesis that $x$ does not affect the variability of $y$. The second equation is essentially the Breusch-Pagan test, except that the variance is calculated under the null that $\beta=0$.

Let $\tilde{y}_{i}=y_{i}-\bar{y}$, $\tilde{x}_{i}=x_{i}-\bar{x}$ and $\tilde{d}_{i}=\tilde{y}_{i}^{2}-\hat{\sigma}^{2}$, where $\hat{\sigma}^{2}=\frac{1}{n}\sum_{i=1}^{n} \tilde{y}_{i}^{2}$. Further, let the $n\times k_{x}$ matrix $\tilde{X}=\left[ \tilde{x}_{i}' \right]$ and the $n$ vectors $\tilde{y}=\left( \tilde{y}_{i} \right)$ and $\tilde{d}=\left( \tilde{d}_{i} \right)$. Then the OLS estimators for $\beta$ and $\delta$ are given by

$$\hat{\beta}=\left( \tilde{X}'\tilde{X} \right)^{-1}\tilde{X}'\tilde{y}$$

$$\hat{\delta}=\left( \tilde{X}'\tilde{X} \right)^{-1}\tilde{X}^{'}\tilde{d} . [9]$$

Let $\theta=\left( \begin{matrix} \beta\\ \delta\end{matrix} \right), \hat{\theta}=\left( \begin{matrix} \hat{\beta} \\ \hat{\delta} \end{matrix} \right)$ and $\hat{\Sigma}=\frac{1}{n}\sum_{i=1}^{n} \left[ \begin{matrix} \tilde{y}_{i}^{2} & \tilde{y}_{i}\tilde{d}_{i} \\ \tilde{y}_{i}\tilde{d}_{i} & \tilde{d}_{i}^{2} \end{matrix} \right]$. The estimator for the variance of $\hat{\theta}$ under the null that $\beta=\delta=0$ and the additional assumption that the conditional skewness and kurtosis of $y_{i}$ do not vary with the values of $x_{i}$, is then given by

$$V\hat{a}r\left( \hat{\theta} \right)={\hat{\Sigma}\otimes\left( \tilde{X}'\tilde{X} \right)}^{-1} . [10]$$

Hence, the score test for $H_{0}:\beta=\delta=0$ or $H_{0}:\theta=0$, is given by

$$S=\hat{\theta}'\left( \hat{\Sigma}^{-1}\otimes\left( \tilde{X}'\tilde{X} \right) \right)\hat{\theta}\underset{\to}{d}\chi_{2k_{x}}^{2} . [11]$$

Additional terms such as the square of a continuous exposure, especially useful for modelling the relationship with outcome variability, can be added as part of $x_{i}$ vector and would be included in both parts of the test. Covariates are regressed out of both the outcome and exposure variables by taking residuals from OLS regression prior to analysis with JLSsc.

In terms of method of moments, let

$$g_{i}=\left( \begin{matrix} \tilde{x}_{i}\tilde{y}_{i} \\ \tilde{x}_{i}\tilde{d}_{i} \end{matrix} \right)=\left( I_{2}\otimes\tilde{x}_{i} \right)\left( \begin{matrix} \tilde{y}_{i} \\ \tilde{d}_{i} \end{matrix} \right) [12]$$

and

$$\hat{V}=\hat{\Sigma}\otimes\left( \tilde{X}'\tilde{X}/n \right) [13]$$

Then, with $\bar{g}=\frac{1}{n}\sum_{i=1}^{n} g_{i}$, it follows that

$$S=n\bar{g}'\hat{V}^{-1}\bar{g}\underset{\to}{d}\chi_{2k_{x}}^{2} . [14]$$

Alternatively, relaxing the constant skewness and kurtosis assumption, a robust version of the test is given by

$$S_{r}= n\bar{g}'\hat{V}_{r}^{-1}\bar{g}\underset{\to}{d}\chi_{2k_{x}}^{2} , [15]$$

where $\hat{V}_{r}=\frac{1}{n}\sum_{i=1}^{n} g_{i}g_{i}'$.

###### Brown-Forsythe methodology

JLSsc can also be set-up using the Brown-Forsythe methodology by replacing $\tilde{d}_{i}=\tilde{y}_{i}^{2}-\hat{\sigma}^{2}$ with $\tilde{d}_{i}=|y_{i}-\overset{̿}{y}|- \frac{1}{n}\sum_{i=1}^{n} |y_{i}-\overset{̿}{y}|$ in [9-15] where $\overset{̿}{y}$ is the median of $y$ (Figure S13). This approach was slightly less conservative in simulations with a skewed residual error (although it did retain correct type I error levels in all scenarios tested) but performed slightly better than the Breusch-Pagan approach in simulations where the exposure was a categorical variable with more than 2 categories (data not shown).

#### Simulation study

The type I error simulations were performed by randomly generating a binary or continuous exposure and testing the associations across all CpG sites in Tsaprouni *et al*.^15^ dataset (data accessible at NCBI GEO database^16^, accession GSE50660). To generate datasets with varying sample size (100, 500, 1000 and 10,000 samples), samples were randomly taken with replacement from the Tsaprouni *et al.* dataset, while adding a small amount of noise by drawing from a random normal distribution with mean zero and the standard deviation set to ten percent of the standard deviation of methylation at that CpG site. The binary and categorical exposures were randomly generated using $Ber(0.5)$ and $N(0,1)$, respectively.

The power simulations were performed using the following model,

$$y_{i}=\alpha+\beta x_{i}+v_{i}+e_{i} ,$$

where $y_{i}$ is the methylation at the randomly selected CpG site for the $i$-th sample and $x_{i}$ is either the binary or continuous exposure ($x_{i} \sim Ber(0.5)$or $x_{i} \sim N(0,1)$). $\alpha$ is the mean methylation from Tsaprouni *et al.* at the randomly selected CpG site ($\mu$), $\beta=\pm w\sigma$ and $v_{i}\sim N(0,zx_{i}\sigma^{2})$, where $\sigma$ is the standard deviation (SD) of methylation from Tsaprouni *et al.* at the CpG site, $w$ is the scaler of the SD for the mean effect and $z$ is the scaler for the variability effect. The direction of $w$ was set to be positive if $\mu\leq0.5$ and negative if $\mu>0.5$. The residual ($e_{i}$) was set to be either normally distributed [$e_{i} \sim N(0,\sigma^{2})$; $w$= (0, 0.1, …, 0.5) and $z$= (0, 0.05, …, 1) for binary exposures and $w$= (0, 0.025, …, 0.3) and $z$= (0, 0.05, …, 0.5) for continuous exposures], heavy-tailed [$e_{i} \sim\sigma t_{4}$; $w$= (0, 0.1, …, 0.5) and $z$= (0, 0.05, …, 1) for binary exposures and $e_{i} \sim\sigma t_{8}$; $w$= (0, 0.05, …, 0.5) and $z$= (0, 0.1, …, 1) for continuous exposures] or skewed [$e_{i} \sim\frac{\sigma}{2}\chi_{1}^{2}$; $w$= (0, 0.025, …, 0.25) and $z$= (0, 0.05, …, 1) for binary exposures and $w$= (0, 0.025, …, 0.3) and $z$= (0, 0.05, …, 0.5) for continuous exposures]. Additional normally distributed simulations were performed using a categorical exposure with three categories [$w$= (0, 0.1, …, 0.5) and $z$= (0, 0.05, …, 1) for the second category and $w$= (0, 0.2, …, 1) and $z$= (0, 0.2, …, 4) for the third category]. Further simulations were performed using the normally distributed residual simulation set for the binary and continuous exposures where a single outlier was simulated to be 5 standard deviations away from $\mu$.

#### Application to offspring gender and gestational age on cord blood DNA methylation

##### Quality control and pre-processing procedures in ARIES

Cord blood samples were collected according to standard procedures. The DNA methylation wet laboratory and pre-processing analyses were performed at the University of Bristol as part of the ARIES project. Following extraction, DNA was bisulphite converted using the Zymo EZ DNA MethylationTM kit (Zymo, Irvine, CA, USA). Following conversion, genome-wide methylation of over 485,000 CpG sites were measured using the Infinium HumanMethylation450 BeadChip according to the standard protocol. The arrays were scanned using an Illumina iScan and initial quality review was assessed using GenomeStudio (version 2011.1).

Samples from all time points in ARIES were distributed across slides using a semi-random approach (sampling criteria were in place to ensure that all time points were represented on each array) to minimize the possibility of confounding by batch effects. In addition, during the data generation process a wide range of batch variables were recorded in a purpose-built laboratory information management system (LIMS). The main batch variable was found to be the bisulphite conversion plate number. Samples were converted in batches of 48 samples and each batch identified by a plate number. The LIMS also reported quality control (QC) metrics from the standard control probes on the 450K BeadChip for each sample. Samples with more than 5% of probes that have detection $p>$ 0.01 were excluded from the analysis. As an additional QC step genotype probes were compared with SNP-chip data from the same individual to identify and remove any sample mismatches. For individuals with no genome-wide SNP data, samples were flagged if there was a sex-mismatch based on X and Y chromosome methylation.

In addition to these QC steps, probes that had detection $p>$ 0.01 for more than 5% of samples were excluded from analysis. After excluding these probes as well as probes on the X or Y chromosomes, a total of 468,611 CpG sites were included in the main analysis. Raw probe intensities were normalized using functional normalization with the meffil package^17, 18^. Methylation levels at each CpG site 5 standard deviations away from the mean were excluded.

##### Gestational age

Gestational age was calculated (in weeks) based on the date of the mother’s last menstrual period when the mother was certain of this, but for uncertain last menstrual periods and conflicts with clinical assessment the ultrasound assessment was used. Where maternal report and ultrasound assessment conflicted, an experienced obstetrician reviewed clinical records and made a best estimate.

### Supplementary Table Legends

Table S1: Characteristics of the mother-offspring pairs with complete covariate information in ARIES.

Table S2: Epigenome-wide association results for gender in cord blood methylation.

Table S3: Epigenome-wide association results for gestational age in cord blood methylation.

### Supplementary Figures

Figure S1: QQ plots for OLS regression (mean test) in type I error simulations. a) binary exposure in 100 samples. b) binary exposure in 500 samples. c) continuous exposure in 100 samples. d) continuous exposure in 500 samples.

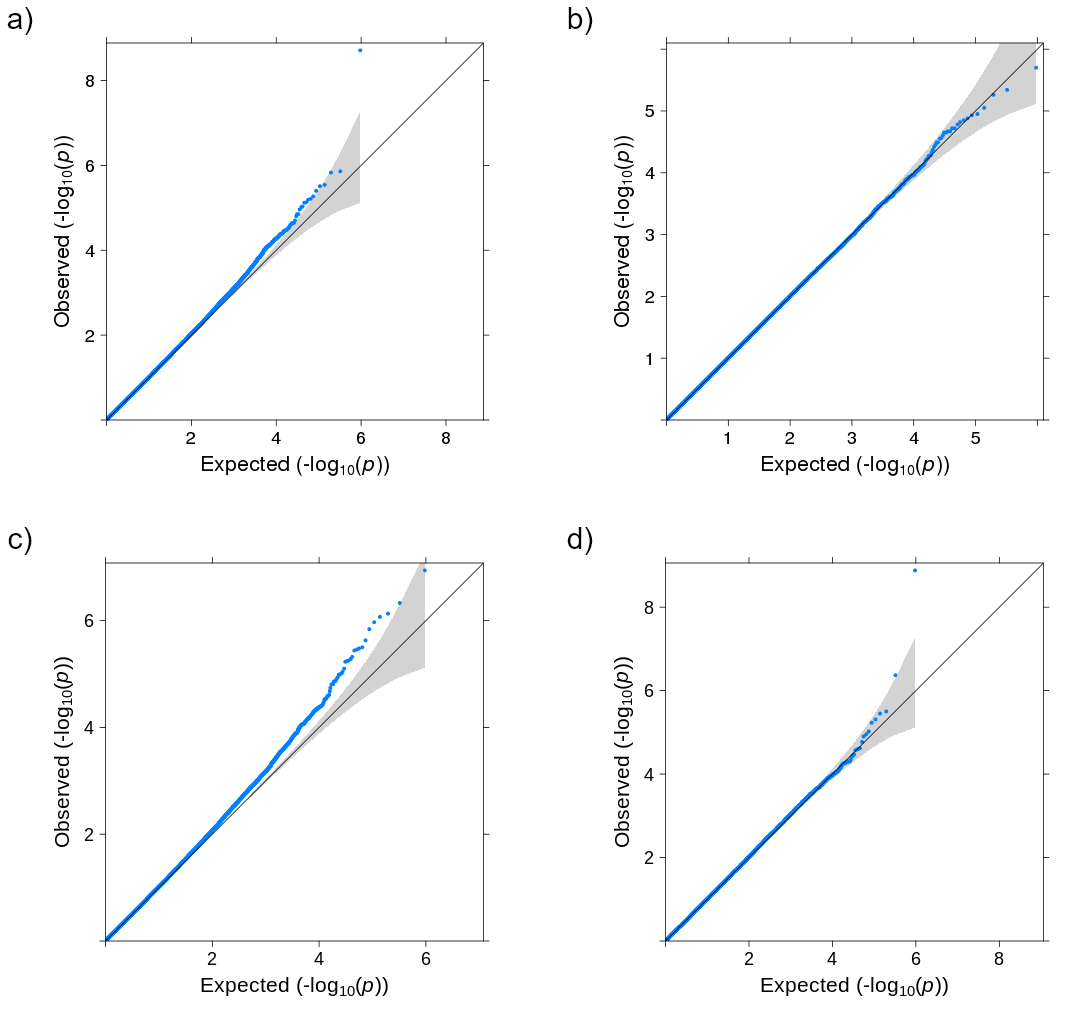

Figure S2: QQ plots for Brown-Forsythe test (variability test) in type I error simulations. a) binary exposure in 100 samples. b) binary exposure in 500 samples. c) continuous exposure in 100 samples. d) continuous exposure in 500 samples.

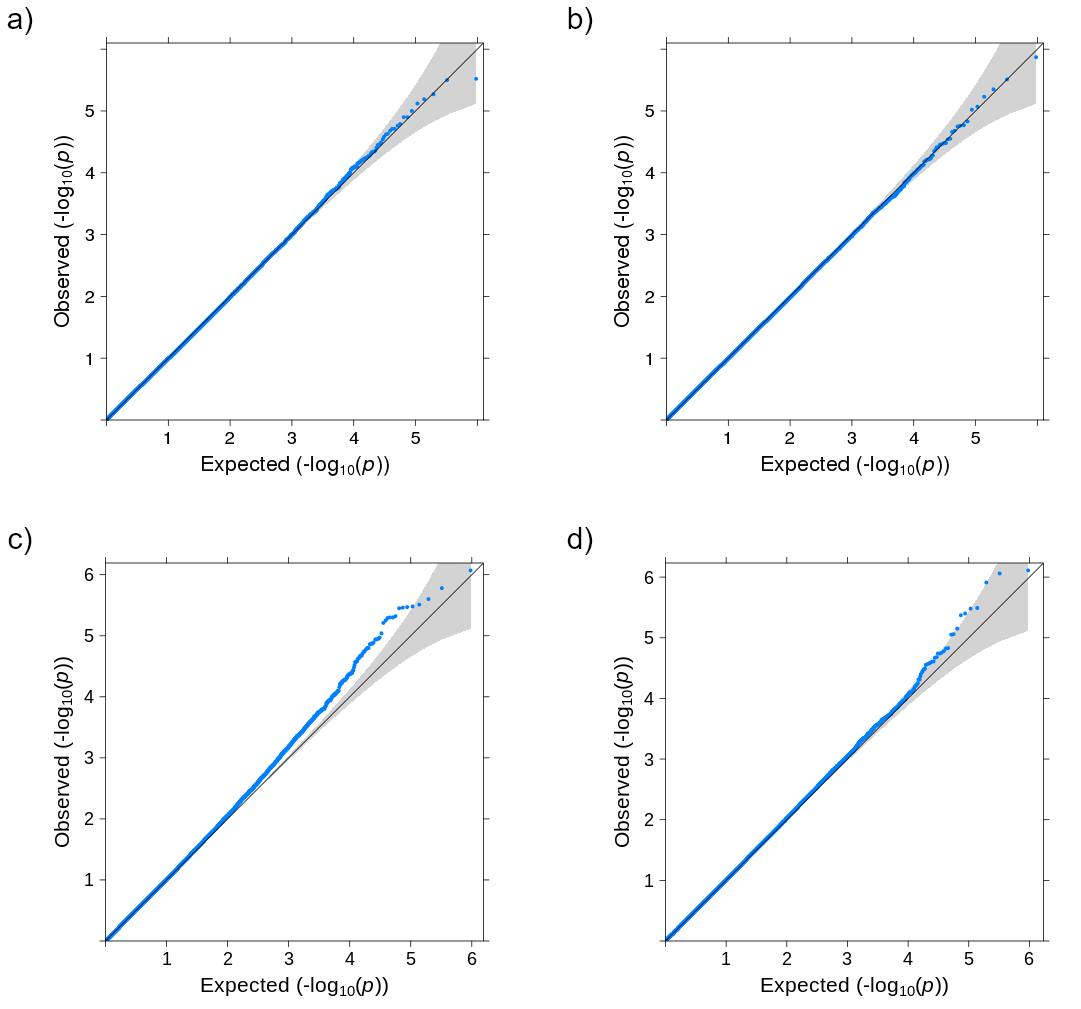

Figure S3: QQ plots for type I error simulations for a binary exposure in 1000 samples. a) Bartlett’s test (variability test). b) LRTv (variability test). c) LRTmv (joint test). d) DGLM (joint test).

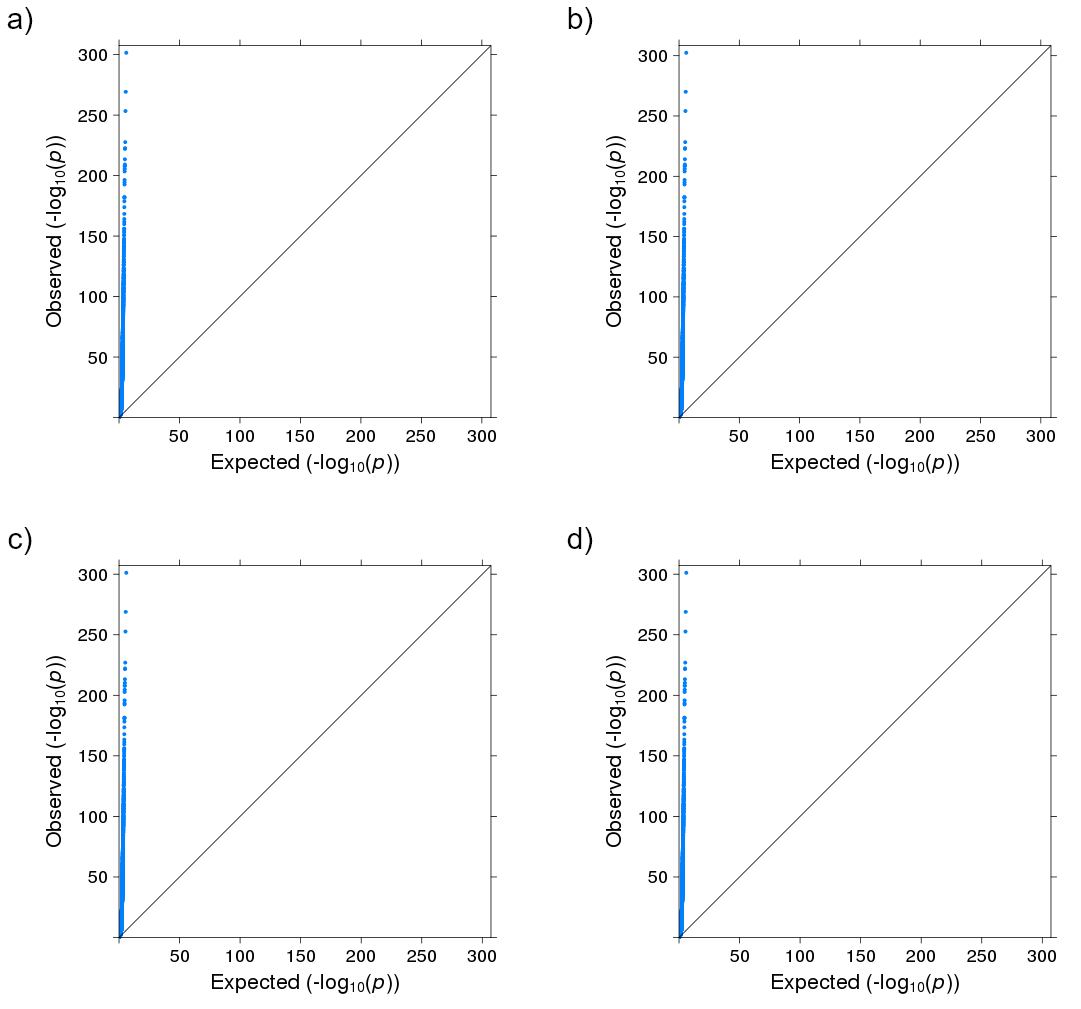

Figure S4: QQ plots for type I error simulations for a binary exposure in 1000 samples after transforming methylation levels using the M-value transformation. a) Bartlett’s test (variability test). b) LRTv (variability test). c) LRTmv (joint test). d) DGLM (joint test).

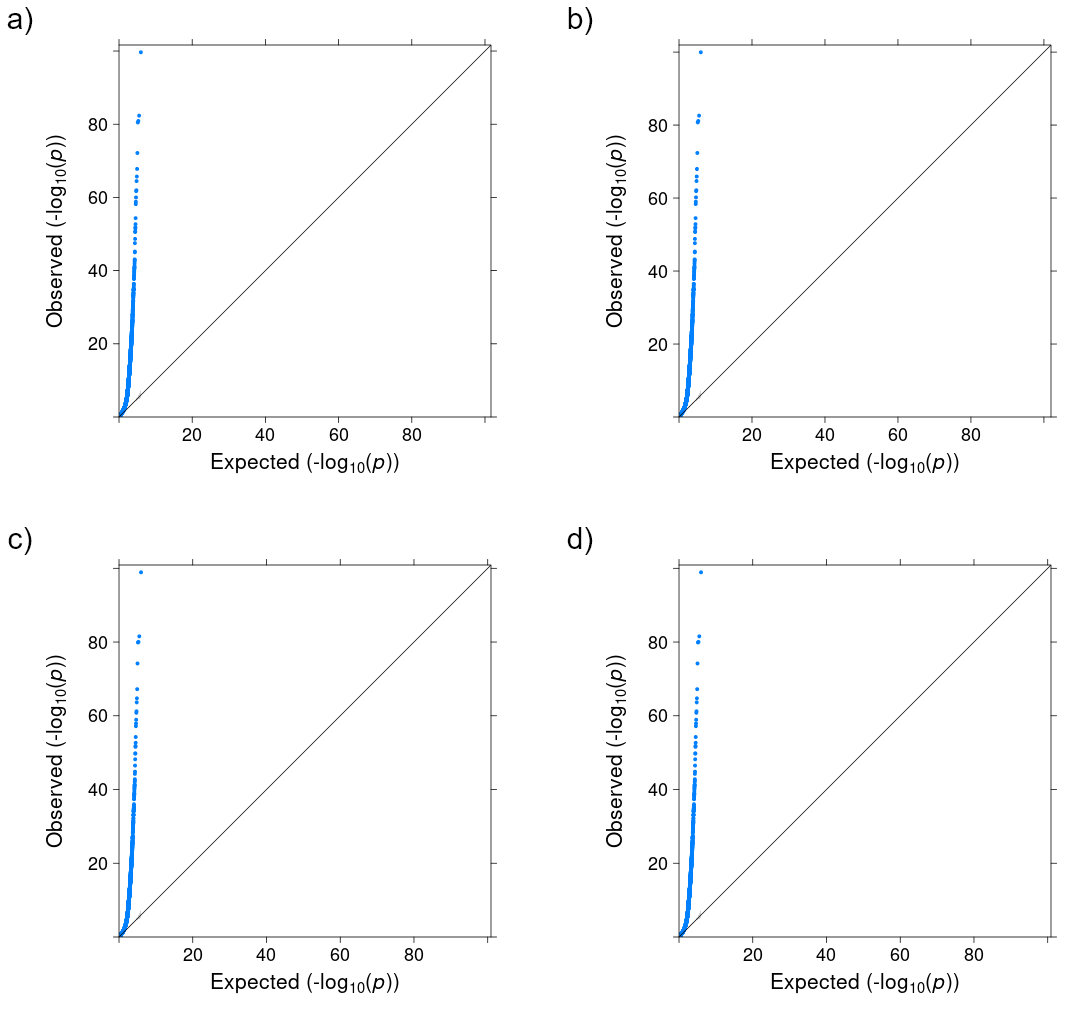

Figure S5: QQ plots for type I error simulations for a binary exposure in 1000 samples after transforming methylation levels using an inverse normal rank transformation. a) Bartlett’s test (variability test). b) LRTv (variability test). c) LRTmv (joint test). d) DGLM (joint test).

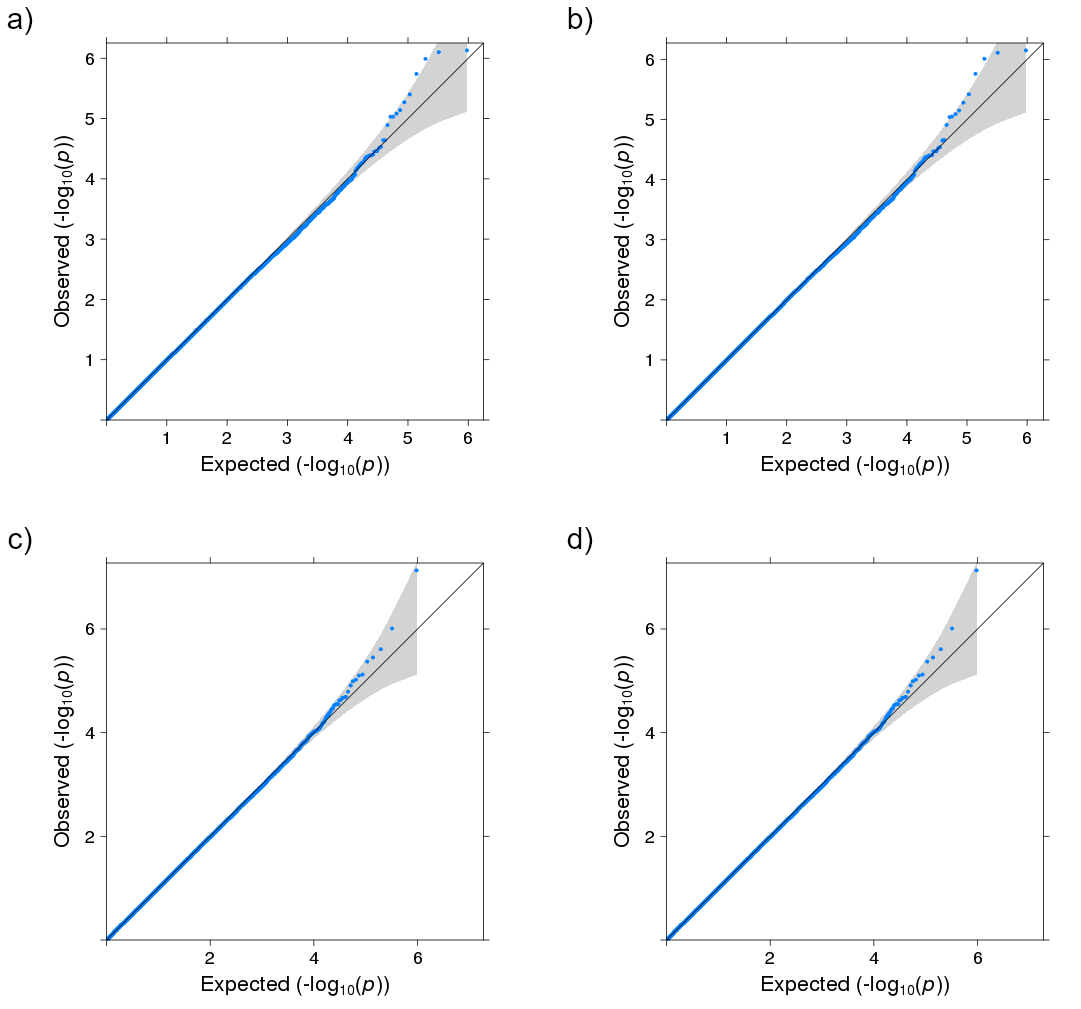

Figure S6: QQ plots for type I error simulations for a binary exposure in 1000 samples after transforming the methylation levels using an inverse normal rank transformation where the residual error is skewed. a) OLS regression (mean test) where there is a variability effect of 0.5 but no mean effect. b) Brown-Forsythe test (variability test) where there is a mean effect of 0.1 but no variance effect.

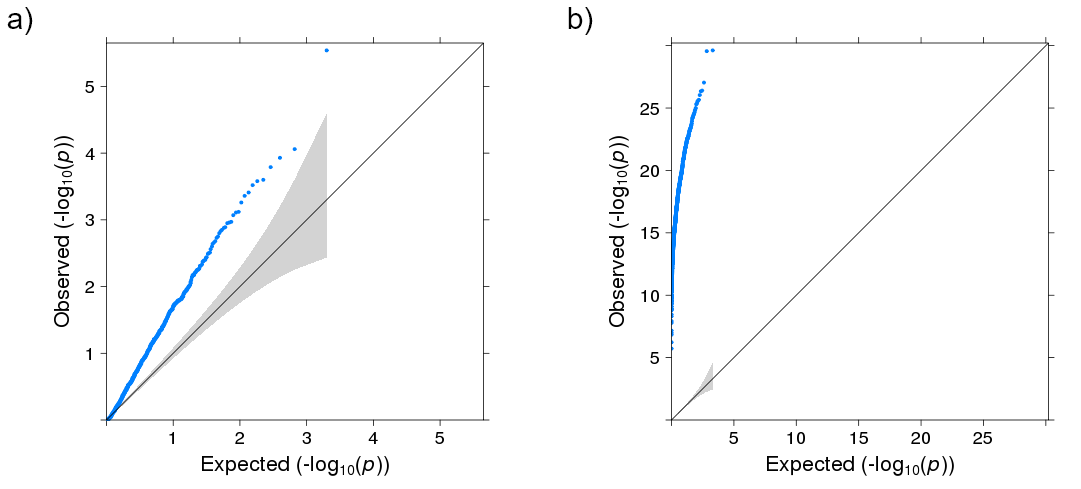

Figure S7: QQ plots for JLSsc (joint test) and JLSp (joint test) in type I error simulations using 1000 samples. a) JLSsc for a binary exposure. b) JLSsc for a continuous exposure. c) JLSp for a binary exposure. d) JLSp for a continuous exposure.

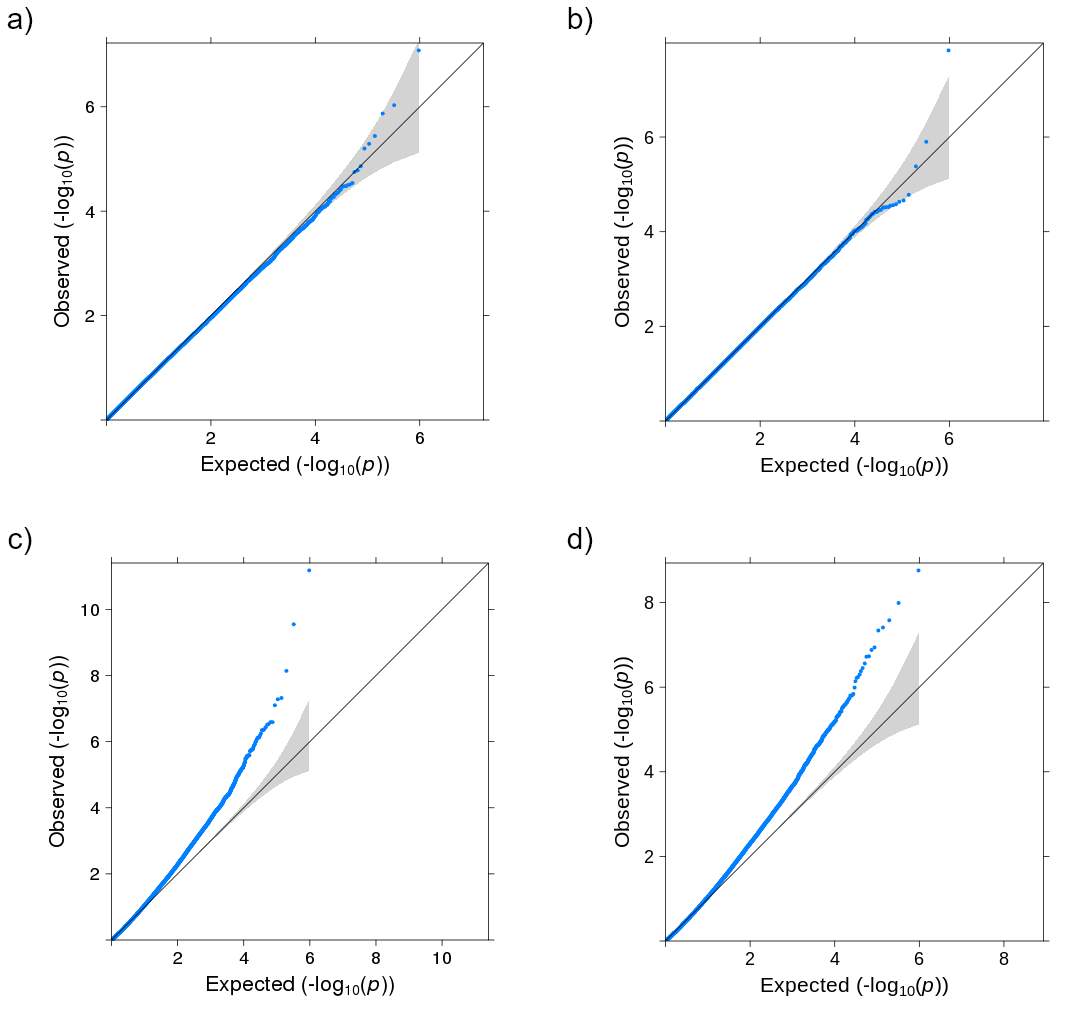

Figure S8: Power simulation results from a heavy-tailed distribution comparing approaches for identifying CpG sites associated with either a mean and/or a variance effect with the exposure at $p<1\times{10}^{-7}$. a) & b) are plots for a binary exposure and c) & d) are plots for a continuous exposure.

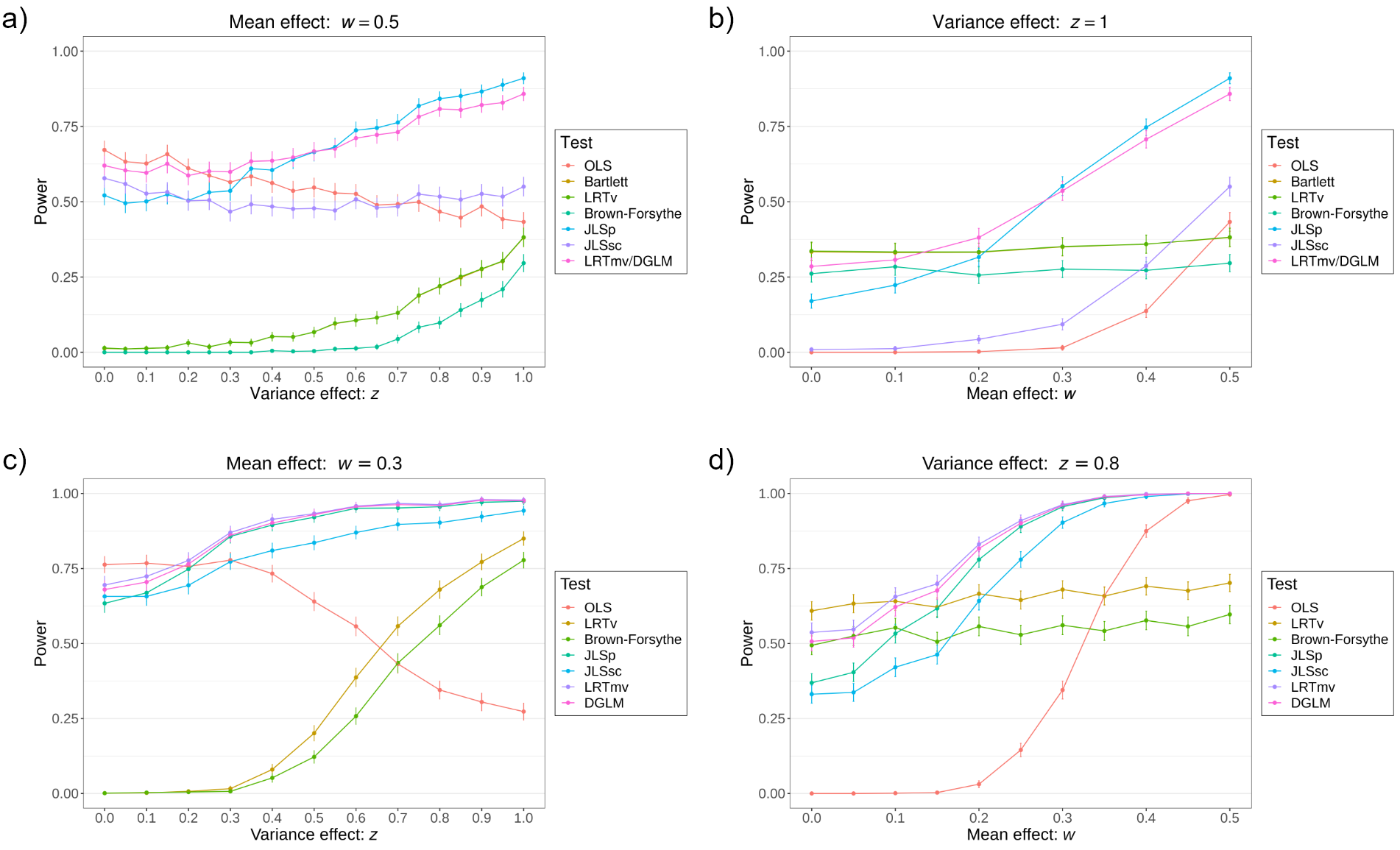

Figure S9: Power simulation results from a skewed distribution comparing approaches for identifying CpG sites associated with either a mean and/or a variance effect with the exposure at $p<1\times{10}^{-7}$. a) & b) are plots for a binary exposure and c) & d) are plots for a continuous exposure.

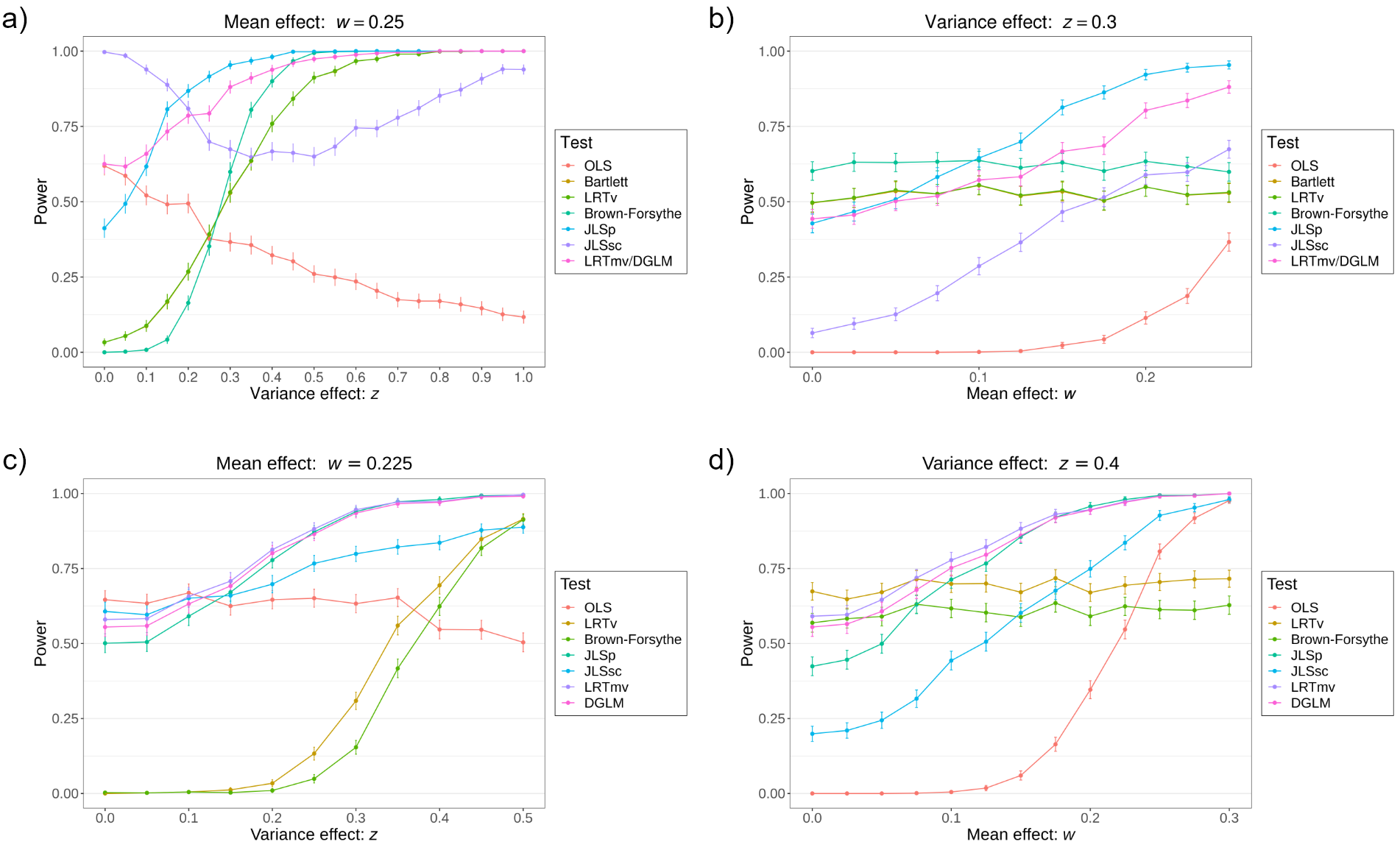

Figure S10: The statistical properties of the JLS approaches (joint tests) for a categorical exposure with 3-levels. a) QQ plot for JLSsc for type I error simulations using 1000 samples. b) QQ plot for JLSp for type I error simulations using 1000 samples. c) Power simulations using 1000 for a mean effect of 0.4. d) Power simulations using 1000 samples for a variance effect of 0.25.

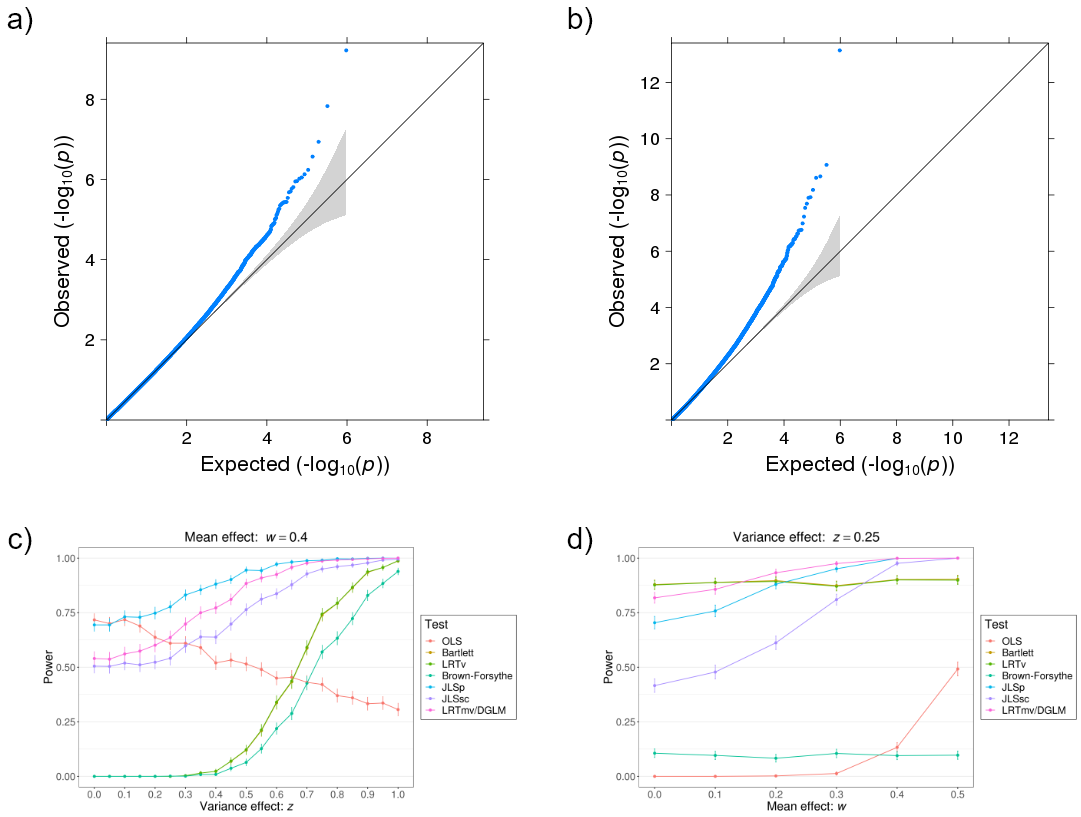

Figure S11: The statistical properties of the JLS approaches (joint tests) in simulations including an exposure squared term for a continuous exposure. a) QQ plot for JLSsc for type I error simulations using 1000 samples. b) QQ plot for JLSsc with an $x^{2}$ term for type I error simulations using 1000 samples (points that were >3×SD were defined as outliers and were removed). c) Power simulations for a mean effect of 0.4. d) Power simulations for a variance effect of 0.7.

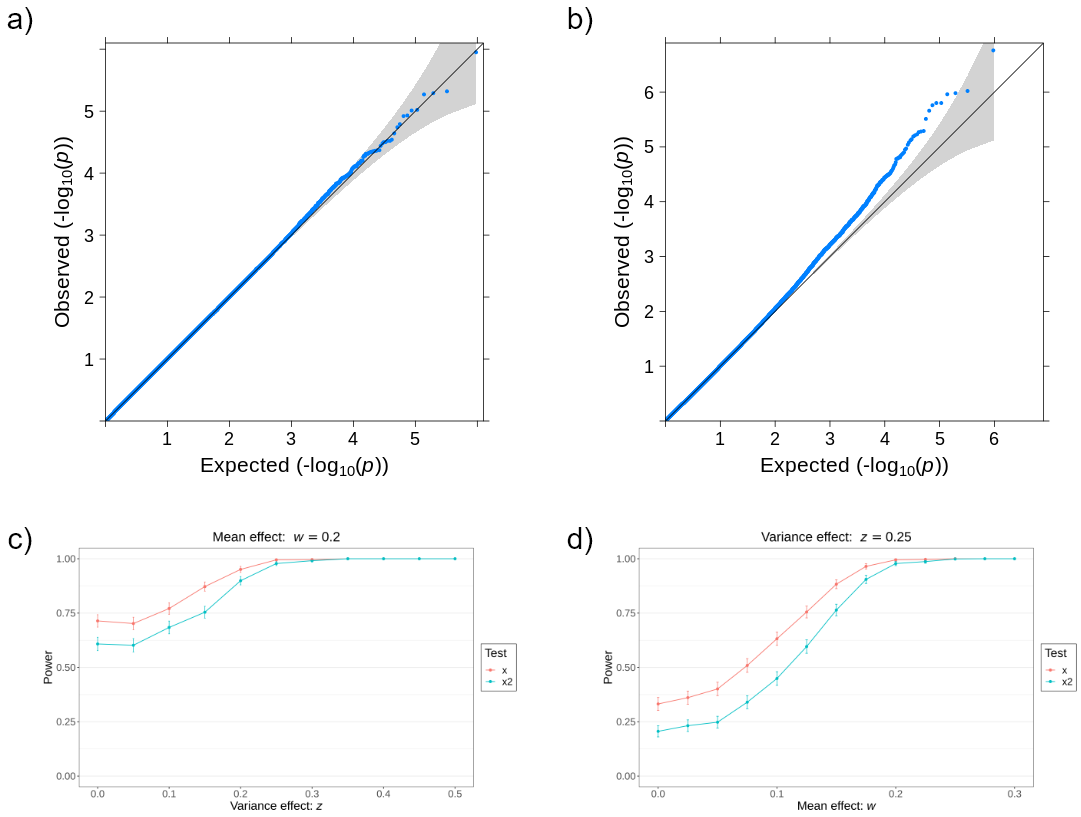

Figure S12: The statistical properties of the JLS approaches (joint tests) in simulations with an outlier for a binary exposure. a) QQ plot for JLSsc for type I error simulations using 1000 samples. b) QQ plot for JLSp for type I error simulations using 1000 samples. c) Power simulations for a mean effect of 0.4. d) Power simulations for a variance effect of 0.7.

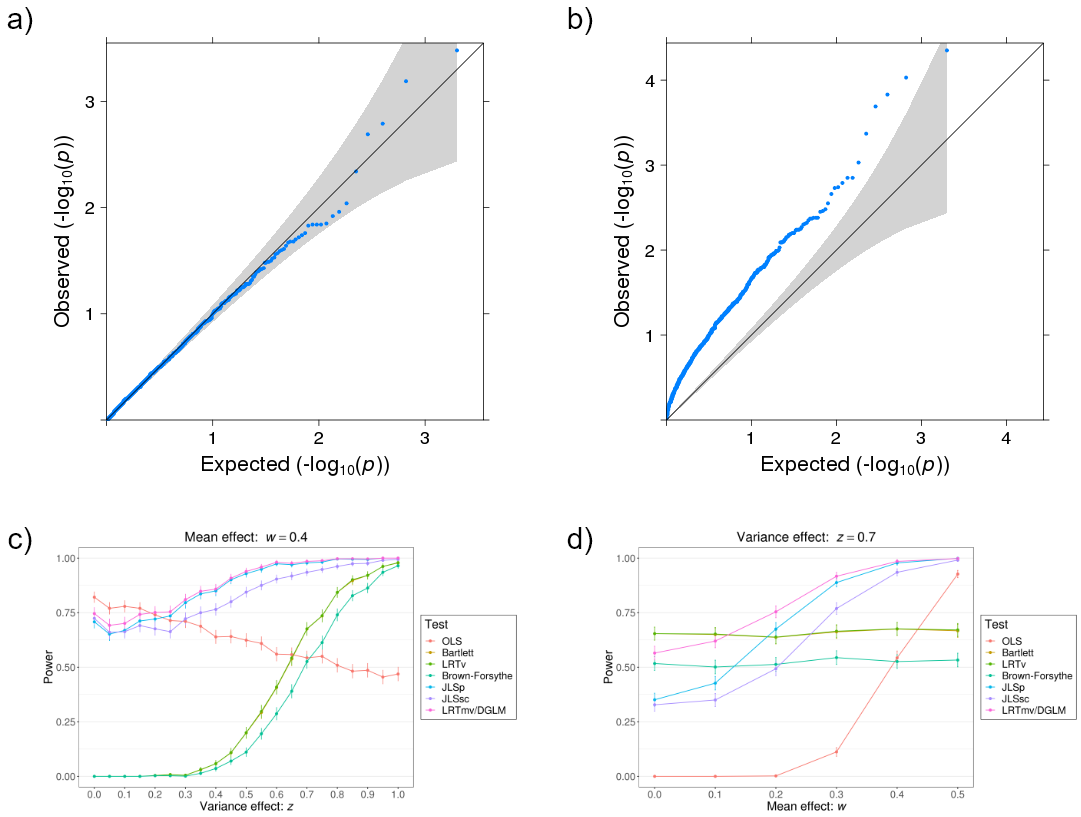

Figure S13: The statistical properties of the JLSsc approach (joint test) using the Brown-Forsythe methodology with a binary exposure. a) QQ plot for JLSsc for type I error simulations using 1000 samples. b) QQ plot for JLSsc based on the Brown-Forsythe methodology for type I error simulations using 1000 samples c) Power simulations for JLSsc using 1000 for a mean effect of 0.4. d) Power simulations for JLSsc using 1000 samples for a variance effect of 0.7.

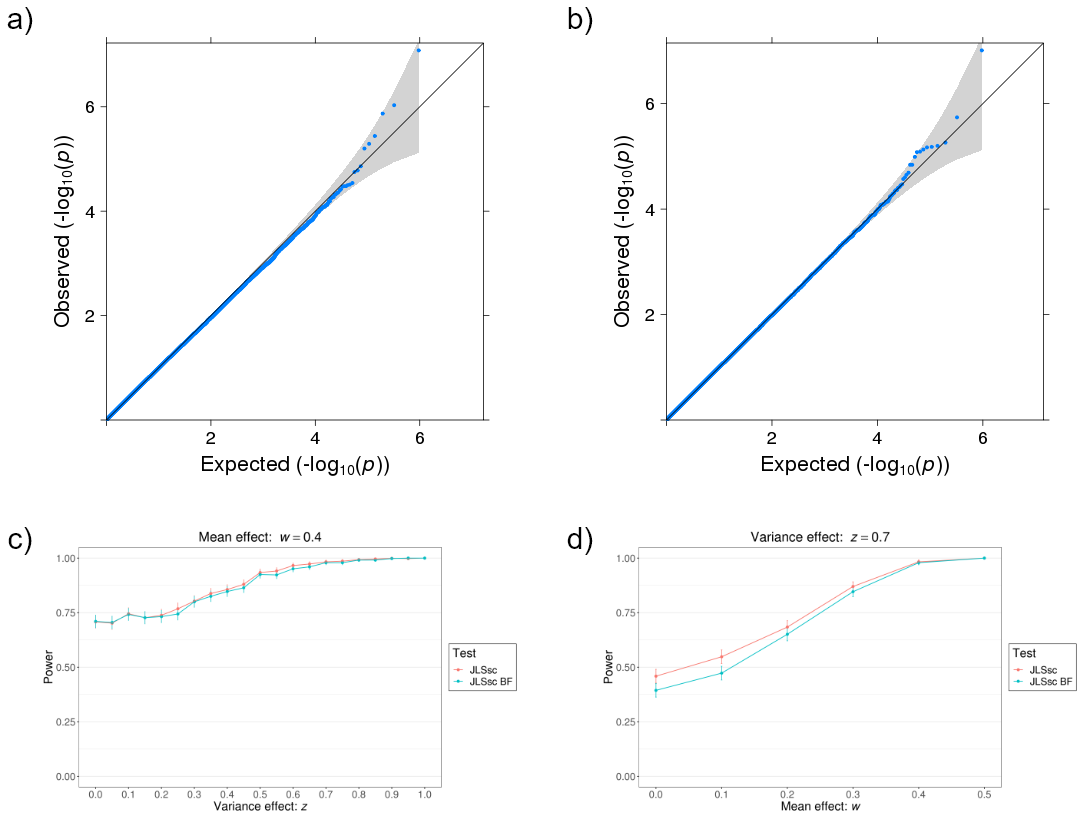
